# A pupillary index of susceptibility to decision biases

**DOI:** 10.1101/247890

**Authors:** Eran Eldar, Valkyrie Felso, Jonathan D. Cohen, Yael Niv

## Abstract

Under what conditions do humans systematically deviate from rational decision making? Here we show that pupillary indices of low neural gain are associated with strong and consistent biases across six different extensively-studied decision making tasks, whereas indices of high gain are associated with weak or absent biases. Lower susceptibility to biases, however, comes at the cost of indecisiveness, or alternatively, prolonged deliberation time. We explain the association between low gain and strong biases as reflecting a broader information integration process that gives greater weight to weak biasing influences. The findings underscore the role of pupil-linked brain states in the generation of decision making biases.

**Significance:** “Framing effects” are demonstrations that people’s decisions can be biased by the way a decision problem is presented, and consequently, people can make decisions that violate the principles of rationality. Using a set of classic decision-making tasks, we show that pupil dilation, previously linked to levels of the neuromodulator norepinephrine and to a tradeoff between narrowly focused and broadly integrative modes of information processing, distinguishes between people who are consistently biased and people who are relatively immune to these effects. Our findings suggest that norepinephrine may drive these individual differences, and that a narrowly focused mode of information processing confers relative immunity to these decision making biases, whereas the integration of a wider range of information results in greater susceptibility.

## Introduction

In certain well-described scenarios, human decision making exhibits systematic deviations from rational behavior. For instance, depending on the precise description of a decision making problem, a particular option could be more or less likely to be chosen even though equivalent information was provided in both descriptions (e.g., “framing effect”; Tversky et al., 1981). These decision biases are typically induced by peripheral aspects of a decision problem, and effects are relatively weak, detectable only when behavior of dozens or even hundreds of participants is averaged (e.g., Levin et al., 1998; Kühberger, 1998). We therefore hypothesized that these biases may predominantly manifest in only a subset of decision makers who process information in a broadly integrative manner, taking into account both central and peripheral aspects of the problem. In contrast, narrowly focusing on a problem’s central aspects may confer relative immunity to these biases. In line with this suggestion, a range of theoretical and experimental work indicates that many decision biases (e.g., framing effects) arise from a process of information integration, and may thus be weaker when the opportunity to integrate information is limited (Usher at al., 2013; Busemeyer et al., 2006; Krajbich & Rangel, 2011, Usher & McClelland, 2004; Busemeyer & Townsend, 1993; Diederich, 1997; Roe et al., 2001; Johnson & Busemeyer, 2005). To test our hypothesis, we focused on pupil diameter measurements that have been suggested to index locus coeruleus-norepinephrine function and neural gain (Servan-Schreiber et al., 1990; Aston-Jones & Cohen, 2005; Joshi et al., 2016), and that we have recently shown to distinguish between people who integrate different aspects of available information versus those who narrowly focus on only the most salient aspect (Eldar et al., 2013, in press). Our goal was to test whether pupil diameter measurements could distinguish between those susceptible to description-induced decision biases and those who were immune to these manipulations, across a variety of well-established tasks.

## Results

We tested human participants on six different decision-making tasks, while measuring their pupil dilation responses during performance of the tasks. Below we first detail the results of each of the tasks, and then provide a mechanistic, computational model that accounts for the findings.

### Anchoring

The role of broad information integration is most obvious in decision biases that arise from the influence of previous events. Such biases essentially reflect integration over time, of present and past information. Perhaps the simplest case is that of the anchoring of estimations to arbitrary values considered in preceding questions (Tversky & Kahneman, 1974). In line with previous studies of anchoring, we asked participants to indicate whether seven different quantities (e.g., the height of the Eiffel tower) were higher or lower than some arbitrary value, and then to estimate the quantities (Jacowitz & Kahneman, 1995). Anchoring was measured by the degree to which a participant’s estimation deviated towards the arbitrary value that the participant was asked to consider initially, as compared to other participants’ estimates. We divided participants into tertiles of low, medium and high mean pupil dilation, and computed the mean anchoring effect for each group. All groups of participants exhibited a significant anchoring effect, regardless of pupillary response (low: *t*_12_ = 3.7, *p* < 0.005; medium: *t*_13_ = 3.3, *p* < 0.01; high: *t*_12_ = 4.2, *p* < 0.005; Fig. 1), with the trend towards stronger anchoring with higher pupillary response (indicating lower gain) not significant (low vs. high: *t*_24_ = 0.81, *p* = 0.43).

**Fig. 1.**
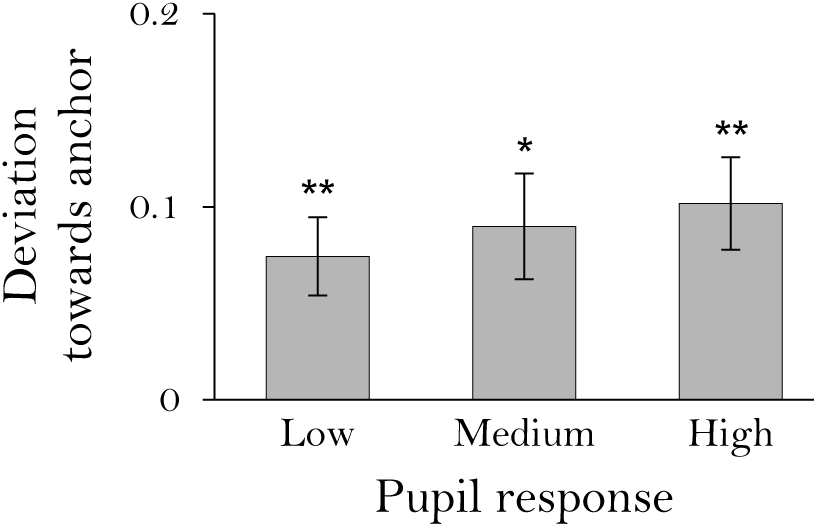
Anchoring. Deviation of participants’ estimates towards the arbitrary anchors which they were asked to consider. Estimates were normalized to the range of 0 to 1. Participants were divided into tertiles based on mean pupillary dilation in response to task stimuli. *n* = 40 participants, *: *p* < 0.01, **: *p* < 0.005, error bars: across-participant s.e.m.

### Persistence of belief

Beliefs formed early in an experiment sometimes persist even in the face of later contradictory evidence (Peterson & DuCharme, 1967). This bias is thought to arise from previously gathered information affecting the weighing of new information (Lord et al., 1979). Thus, this bias too may depend on integration of information over time. To test persistence of belief, we presented participants with a series of colored balls while asking them to judge from which of two urns the balls were more likely to have come. The two urns differed in the proportion of balls of each color, and thus, in the probability of being the source of the series of balls (Fig. 2A). The order of the balls presented was predetermined so as to initially favor one urn (first 30 balls), and then the other (last 60 balls). We quantified persistence of belief bias by the degree to which participants continued to choose the initially-favored urn during the second part of the sequence. In this experiment, only participants with high pupillary responses showed significant persistence of belief (*t*_11_ = 2.8, *p* < 0.05). In contrast, participants with low pupillary responses updated their estimates similarly in the first and second part of the experiment (*t*_22_ = 2.4, *p* < 0.05; Fig. 2B).

**Fig. 2.**
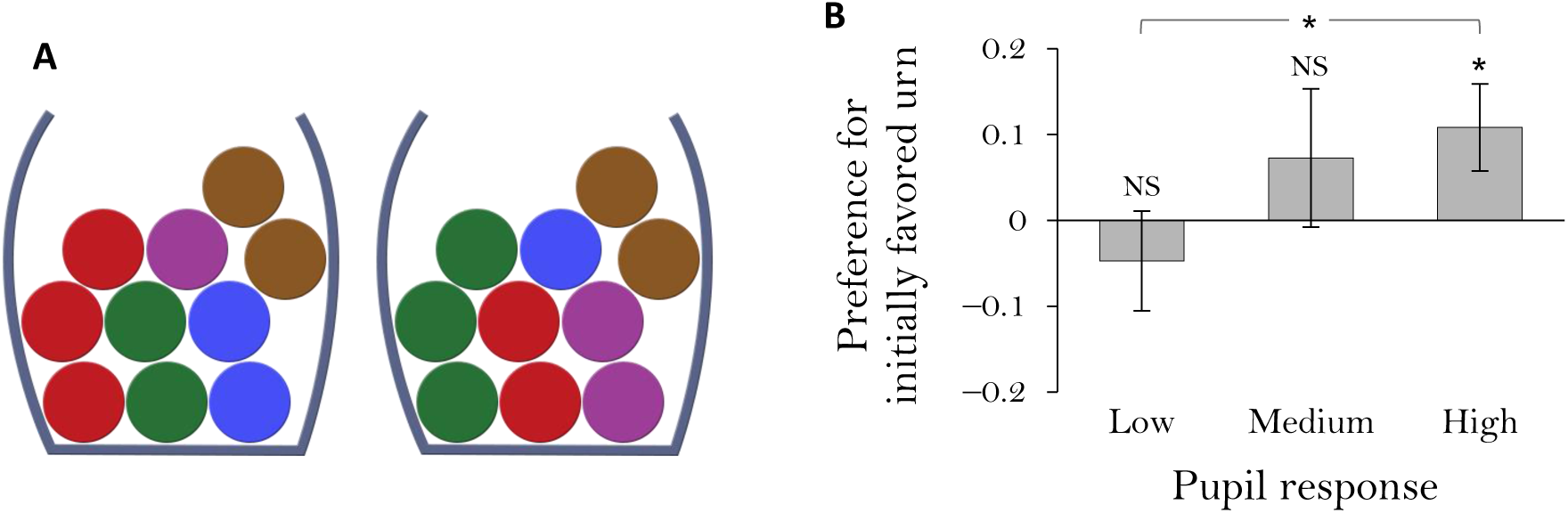
Persistence of belief. (A) The two urns presented to participants contained different proportions of balls of different colors. Balls drawn were pre-determined such that at first it would seem that they were drawn from one urn, whereas later evidence would suggest the other urn. (B) Preference of the initially favored urn during the last 60 balls (which were consistent with the other urn). Preferences were indicated on a scale between -1 and 1. An optimal observer would be indifferent on average. *n* = 35 participants, NS: *p* > 0.1, *: *p* < 0.05, error bars: across-participant s.e.m.

### Attribute framing

Many decision biases arise not from the temporal structure of the problem, but rather from integration of its multiple attributes. We tested four types of biases that are found in multi-attribute multi-alternative decision problems: three types of framing effects and one instance of sample-size neglect. In the first, attribute framing task, participants evaluated items of three different types (ground beef, student exam performance and gambles), whose attributes were framed either positively or negatively (Levin et al., 1985). For example, student exam performance could be described in terms of % correct (positive frame) or % incorrect (negative frame). Biased evaluations (i.e., significantly higher evaluations for positively framed items as compared to those framed negatively) were seen only in participants with high pupillary responses (indicating low gain; *t*_13_ = 2.17, *p* < 0.05; Fig. 3).

**Fig. 3.**
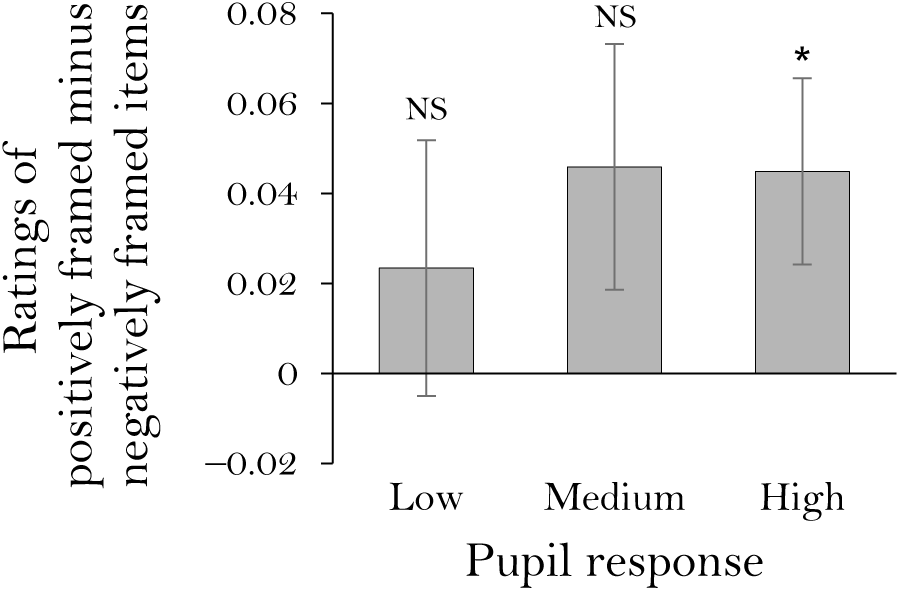
Attribute framing. Difference in evaluation of items framed positively versus negatively. Items were rated on a scale between 0 and 1. Positive values indicate higher evaluations for items framed positively. *n* = 43 participants, NS: *p* > 0.1, *: *p* < 0.05, error bars: across-participant s.e.m.

### Risky choice framing

In the second framing task, participants chose between a certain and an uncertain outcome, both framed either as gains or as losses (Tversky et al., 1981; Van Schie & Van Der Pligt, 1995). For example, the outcome of a treatment program could be described as’200 people (out of 600) will be saved’ or as’400 people (out of 600) will die’. This manipulation exploits people’s tendency to be risk averse in the domain of gains, but risk seeking in the domain of losses (Kahneman & Tversky, 1979). Here again, only participants with high pupillary responses exhibited the effect of framing, in this case, greater risk aversion when outcomes were framed in terms of gains rather than losses (t_13_ = 2.34, *p* < 0.05; Fig. 4).

**Fig. 4.**
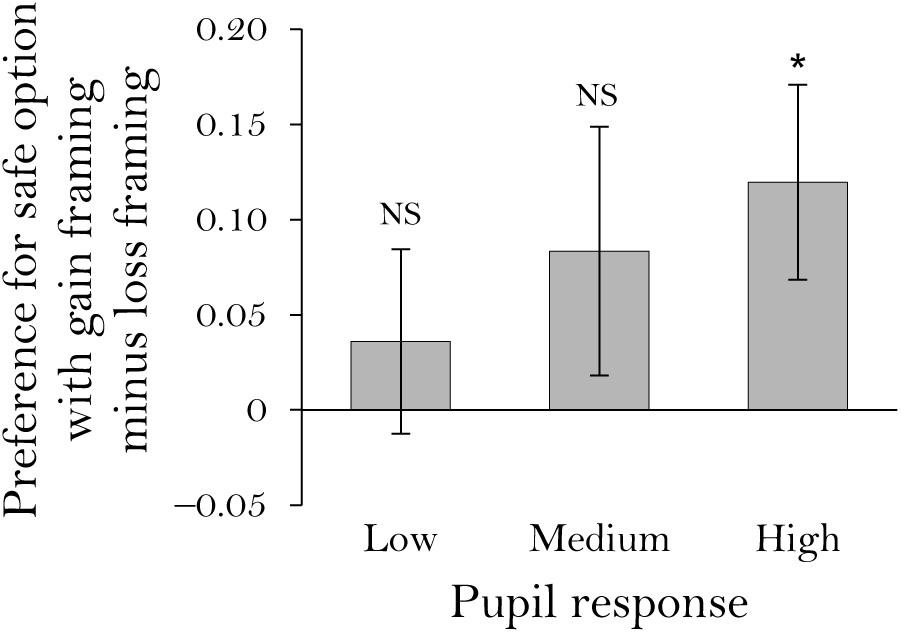
Risky choice framing. Increase in risk aversion when outcomes were described in terms of gains as opposed to losses. Preferences were indicated on a scale between -1 and 1. *n* = 42 participants, NS: *p* > 0.2, *: *p* < 0.05, error bars: across-participant s.e.m.

### Task framing

In the third framing task, participants were asked to either accept or reject one of two options. One ‘enriched’ option was described on more positive as well as more negative dimensions than the other, ‘impoverished’, option (Shafir, 1993). People have been shown to be biased towards the enriched option regardless of whether they are accepting it or rejecting it, presumably because the enriched option provides more reasons to do either. Here, participants with medium or high pupillary responses, but not with low pupillary responses, showed a significant preference for the enriched option across both “accept” and “reject” tasks (medium: *t*_13_ = 3.5, *p* < 0.005; high: *t*_13_ = 2.2, *p* < 0.05; Fig. 5).

**Fig. 5.**
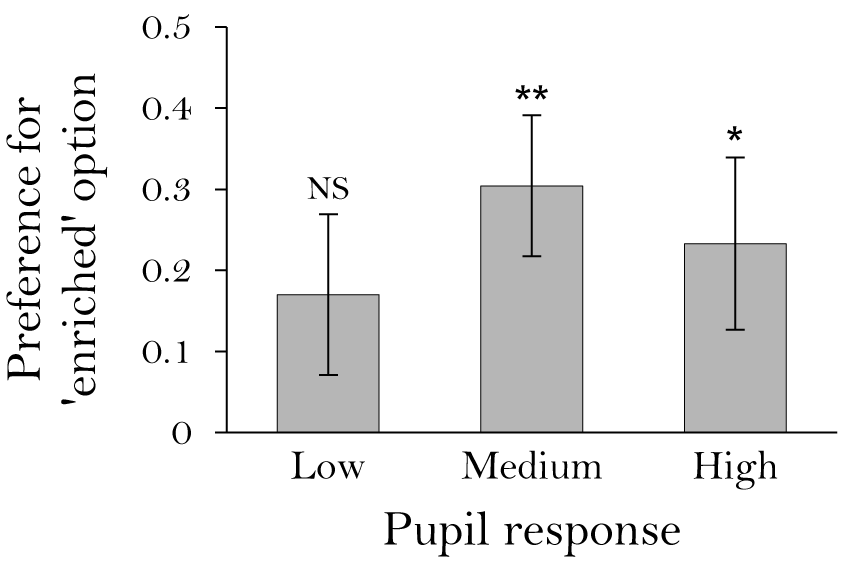
Task framing. Preference to both accept and reject the enriched option more than the impoverished option. Preferences were indicated on a scale between -1 and 1. *n* = 42 participants, NS: *p* > 0.1, *: *p* < 0.05, **: *p* < 0.005, error bars: across-participant s.e.m.

### Sample-size neglect

A different bias that involves multi-attribute integration can be seen when people try to determine, based on the results of several coin tosses, whether the coin is biased to heads or tails. Deciding on the direction of the bias is trivial (i.e., by choosing the more frequent outcome), but people’s certainty about this direction is typically determined solely by the ratio between heads and tails (Griffin & Tversky, 1992), which neglects to account for the sample size of the evidence (i.e., 10 to 6 should induce greater certainty than 5 to 3). An optimal judge would instead simply rely on the difference between heads and tails (see Methods for details). Ratios such as 9 to 7 or 6 to 5 are unlikely to be computed precisely, and thus, relying on ratios necessitates a decision process that integrates both raw numbers. In contrast, the difference between the numbers, which suffices for an optimal decision, can easily be computed precisely, thereby reducing the evidence to a single attribute and avoiding the need for integration.

We asked participants how certain they were that a coin was biased in favor of heads given different sets of outcomes. Participants with medium and high pupil responses exhibited significant sample-size neglect (medium: *t*_12_ = 6.3, *p* < 10-5; high: *t*_11_ = 4.1, *p* < 0.005), whereas those with low pupil responses exhibited only a trend-level effect (*t*_11_ = 2.1, *p* = 0.06) that was significantly weaker than in the other groups (vs. medium: *t*_23_ = 2.5, *p* < 0.05; vs. high: *t*_22_ = 2.6, *p* < 0.05; Fig. 6). Accordingly, responses of participants with low pupil responses reflected more precise inferences than responses in the other groups (vs. medium: *t*_23_ = 2.7, *p* < 0.05; vs. high: *t*_22_ = 3.5, *p* < 0.005). In addition, the *difference* between heads and tails predicted the estimates of participants with low pupil responses better than the ratio between heads and tails (beta difference 0.56 ±0.13, *t*_11_ = 4.3, *p* < 0.005), but this was not true for the estimates of participants with medium (beta difference 0.10 ±0.09, *t*_12_ = 1.1, *p* = 0.31; difference from low group: *t*_23_ = 3.0, *p* < 0.01) and high (beta difference -0.04 ±0.14, *t*_11_ = -0.2, *p* = 0.81; difference from low group: *t*_22_ = 3.1, *p* < 0.005) pupil responses, indicating that only low-pupil-response participants primarily relied on the difference, not the ratio, between heads and tails.

**Fig. 6.**
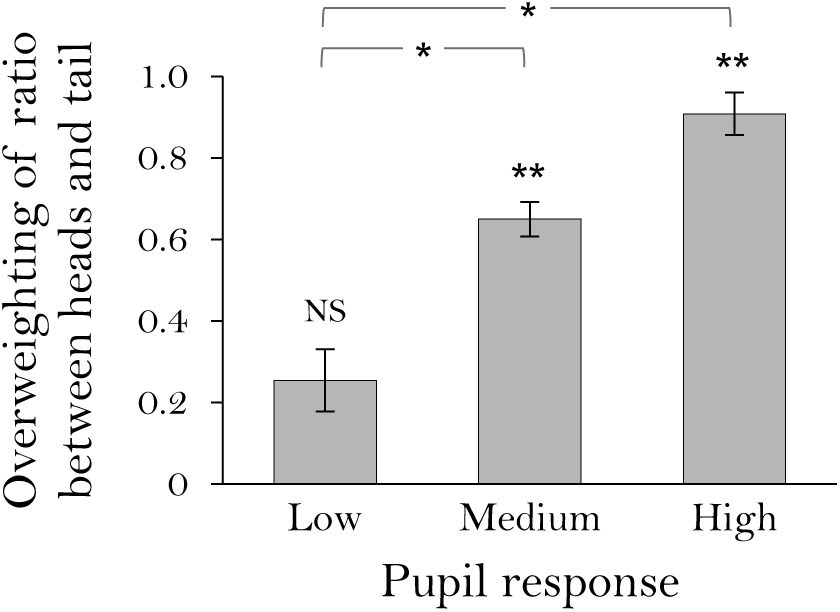
Sample-size neglect. Measured as the overweighting of the ratio between heads and tails relative to the weight given to the optimal inferences (see Methods). *n* = 37 participants, NS: *p* > 0.05, *: *p* < 0.05, **: *p* < 0.005, error bars: across-participant s.e.m.

### Overall susceptibility to biases

The high consistency across tasks of differences in decisionmaking biases between participants with low and high pupil responses is striking – in each one of the six experiments, participants with high pupil responses were more strongly biased. Thus, we compared the average normalized bias effect across all tasks between the groups. Across tasks, participants with low pupillary responses showed significantly weaker biases than participants with high (p < 0.0005, permutation test) and medium (p < 0.01) pupillary responses (Fig. 7). Similarly, the average across-participant correlation between pupillary response and bias effect was significantly greater than zero (mean *r*_s_ = 0.17, *p* < 0.05, permutation test). These results suggest that pupillary response indexed general susceptibility to decision making biases in our experiments.

**Fig. 7.**
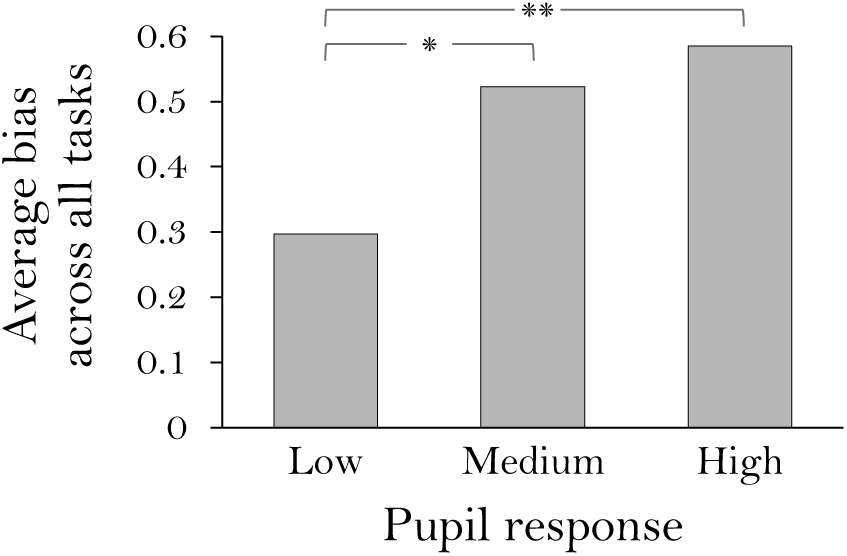
Overall susceptibility to biases. Average normalized bias effect across all experiments. The values 0 and 1 correspond to the minimal and maximal biases exhibited by any subset of 14 participants in each experiment. *n* = 44 participants, *: *p* < 0.01, **: *p* < 0.0005, permutation test.

### Computational model

We next used Usher & McClelland’s (2004) decision model (Fig. 8A) to illustrate how a participant’s pupil response may be related to the formation of decision biases such as those induced by framing. A wide range of evidence indicates that attenuated pupil responses, which are anti-correlated with baseline pupil diameter, reflect a high level of sustained norepinephrine release (Aston-Jones & Cohen, 2005; Eldar et al., 2013; Joshi et al., 2016). Norepinephrine is thought to increase neural gain–that is, to strengthen the impact of all inputs on neural responses (Servan-Schreiber et al., 1990)–an effect that has been used to explain behaviors associated with reduced pupil response in a variety of information processing tasks (Eldar et al., 2013, in press). Thus, we simulated the effect of neural gain on a framing bias in a multi-alternative, multi-attribute decision problem.

Consider a choice between two vacation destinations: Destination A has gorgeous beaches and coral reefs but a high petty crime rate, whereas Destination B has average beaches and an average crime rate. The model assumes that on each time step, attention selects one of the attributes (i.e., ‘beaches’ or ‘crime rate’) at random and the evidence it provides is accumulated at the decision layer. Since framing effects are typically conceptualized as attentional biases that exert their effect during the decision process (Levin et al., 1998), they can be naturally implemented in the model as a tendency to select certain attributes more often than others. For instance, framing the question as to where *to go* would direct attention to more frequently select the positive attribute (beaches), whereas framing the question as where *not to go* would draw attention towards the negative attribute (crime rate).

If decision making requires many time steps, even a small bias accumulates and can determine the result of the decision process. In contrast, if the decision requires only a small number of time steps, the systematic effect of the bias would be minimized. This is where neural gain comes into the picture: lower gain diminishes the effect of evidence on the decision process, increasing the number of time steps required to reach a decision. As a result, biases are stronger (Fig. 8B). In this way, changed in neural gain can explain the robust association of framing effects with high pupil dilation (indicative of low gain) reported above. We note that this account does not rely on a specific decision model – similar results will be obtained by any decision model that involves gradual integration of information.

**Fig. 8.**
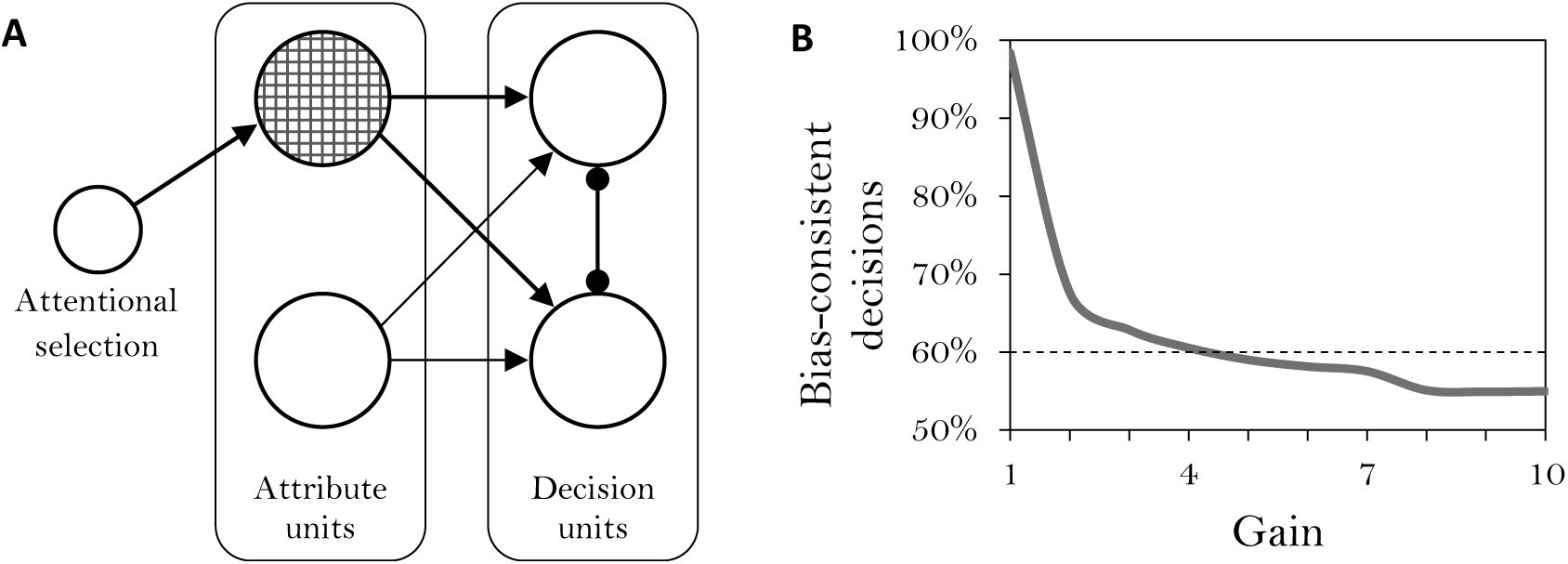
An illustration of the effect of gain on the manifestation of a decision bias. (A) An information accumulation model. On each time step, one attribute is randomly selected, and the evidence in favor of each item is accumulated by the competing decision units. Bias was implemented as a tendency to select one of the attributes more frequently. (B) Proportion of bias-consistent decisions made by the model as a function of gain. With higher gain decisions are less biased. The decision process was simulated 100,000 times with each level of gain. The dashed line indicates the proportion that would be needed to detect a statistically-significant bias given a sample of 100 decisions (binomial test).

### The cost of weaker biases

While high gain is associated with weaker biases, this comes at the cost of limited integration, leading to potential decisions that are based on fewer samples and are thus less certain to be correct. Uncertainty concerning a potential decision is thought to affect the likelihood of executing the decision (Daw et al., 2005), and thus we may expect that high gain would be associated with a higher rate of indecision (e.g., expressing indifference between available options). Indeed, in the multi-alternative decision problems, participants with low pupil responses (indicative of high gain) chose not to decide more often than participants with high pupil responses (Fig. 9A; *p* < 0.05, permutation test).

Alternatively, failure to reach a definitive, integrative decision might induce a deliberative, time-consuming compensatory process whose purpose is to increase decision certainty (Glöckner & Betsch, 2008; Daw et al., 2005). In line with this view, those participants with low pupillary responses who avoided indecision took more time to make decisions relative to those who did not avoid indecision (*p* < 0.05, permutation test), and to those with stronger pupillary responses (*p* < 0.005, Fig. 9B). Moreover, taking more time seems to have restored some level of information integration, since in participants with low pupillary responses longer response times were associated with stronger biases (mean *r*_s_ = 0.30, *p* < 0.05, permutation test; Fig. 9C).

**Fig. 9.**
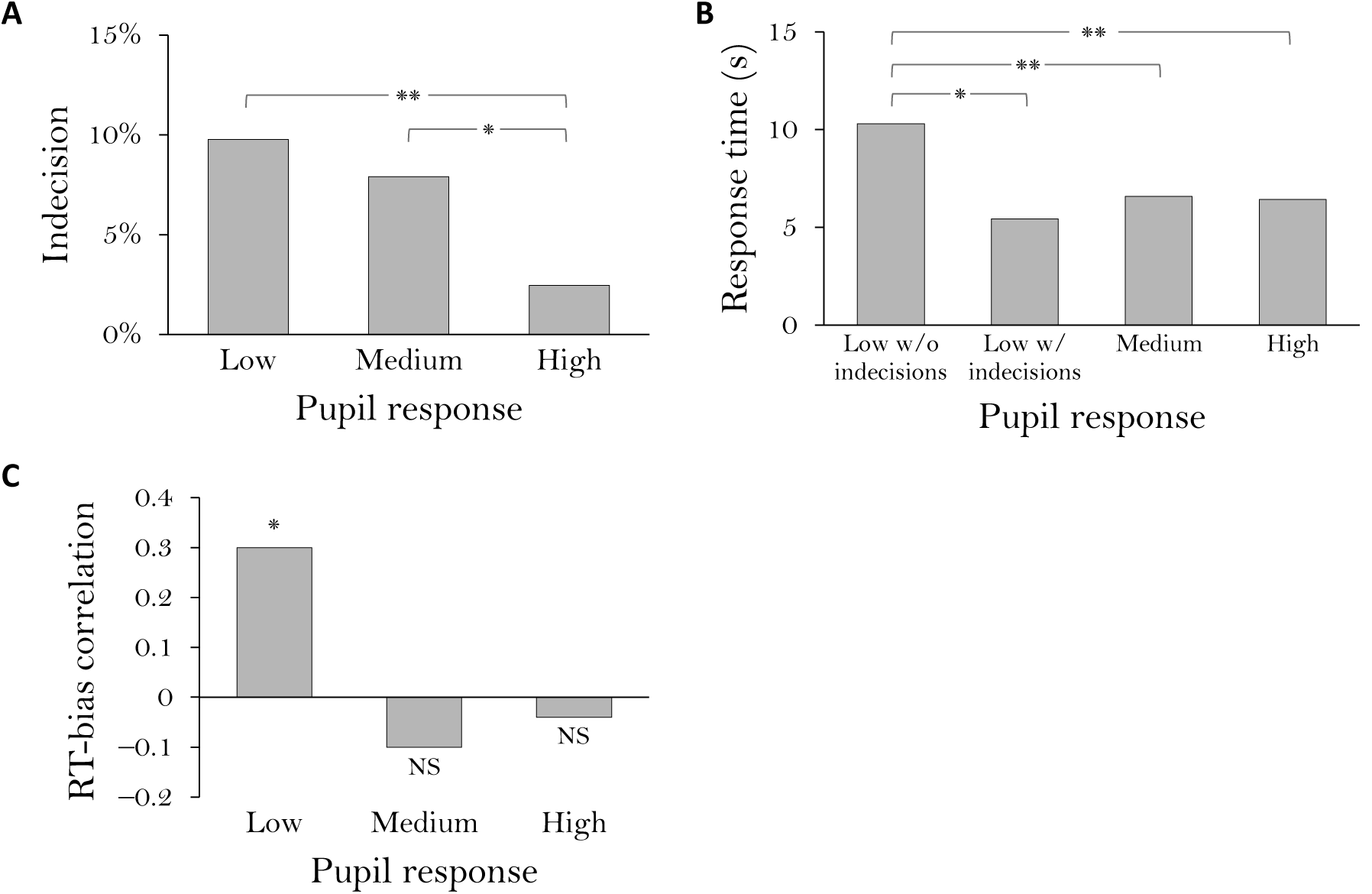
Indecision and response time (RT). (A) Proportion of choices expressing indifference between available options. Data include tasks that involved choice between two alternatives, where participants could choose not to favor either option (i.e., risky choice framing, task framing and persistence of belief tasks). (B) Response time in two-alternative decision problems in which decision time was measureable (i.e., risky choice framing, task framing). The low pupil response group was further divided into participants who exhibited indecisions and those who did not. (C) Mean across-participant Spearman correlation between average response time and bias effect size. Data include tasks in which the source of bias was continuously present (i.e., the framing and sample-size neglect tasks). *n* = 44 participants, NS: *p* > 0.2, *: *p* < 0.05, **: *p* < 0.005, permutation test.

## Discussion

We found an association between low pupil responses, which are thought to indicate high neural gain, and weaker biases across six different tasks that involve integration of multiple pieces of information. However, weaker biases came at the cost of indecisiveness or, alternatively, longer deliberation time. Conversely, participants with high pupil responses showed remarkably consistent biases–dozens or even hundreds of participants are typically needed to demonstrate a decision making bias, whereas we show that when selected for high pupil responses, a small group of at most 15 participants consistently exhibited statistically-significant biases across six different tasks.

It can be argued that the *weaker* biases we found in participants with low pupil responses simply reflect stronger effort by these participants to answer the questions optimally. However, low pupil responses are widely used as indicators of weaker, not stronger effort (Kahneman, 1973). Furthermore, high incentives to answer correctly, which presumably increase effort, do not weaken decision biases such as those that are evoked by framing (Tversky & Kahneman, 1981). In addition, we have previously shown that low and high pupil responses are associated with similar levels of performance in a reward learning task (Eldar et al., 2013) and in a word recognition memory task (Eldar et al., in press).

In this work we considered the view that decision biases emerge from a process of information integration. This view is supported by a variety of experimental and theoretical findings (Usher at al., 2013; Busemeyer et al., 2006). While biases could reflect the operation of other mechanisms, our findings lend further support to the role of information integration in a diverse set of decision making biases, by showing that susceptibility to biases is tracked by a pupillary index of gain, previously linked to behavioral and neural markers of integration (Eldar et al., 2013s; in press).

Our findings suggest that high gain is associated with more optimal decisions. However, the tasks we used – drawn from classic studies of decision making–were specifically designed to be sensitive to people’s tendency to integrate irrelevant biasing cues into the decision process. It is possible, however, that in more complex, real-life decisions that demand consideration of a broad range of factors (e.g., choosing a job or house; Usher et al., 2011; Rusou et al., 2013) the broader integration that low gain permits may confer benefits for decision making.

## Materials and Methods

### Participants

44 Princeton University students (mean age 19.5, age range 18-23, 28 females) performed the experiment. Participants gave written informed consent before taking part in the study, which was approved by the university’s institutional review board. Participants received course credit for participation.

### Stimuli

Stimuli were generated using the Processing programming environment (Reas & Fry, 2007). To minimize luminance-related changes in pupil diameter, stimuli were made isoluminant with the background by adjusting their colors using the flicker-fusion procedure (Lambert et al., 2003) on the display system that was used in the experiment. Stimuli were presented on a computer screen using MATLAB software (MathWorks) and the Psychophysics Toolbox (Brainard, 1997).

### Anchoring experiment

Participants answered two questions about each of 7 quantities (e.g., the height of the Eiffel tower). They first indicated whether the quantity was greater or less than an anchor value. Next, they estimated the quantity. Each quantity was coupled with a low anchor for half of the participants and with a high anchor for the other half. Each participant was presented with a low anchor for half (3 or 4) of the quantities, and with a high anchor for the other half. Quantities and calibrated anchor values were taken from a previous study (Jacowitz & Kahneman, 1995), including: length of the Mississippi river, population of Chicago, number of babies born per day in the US, height of mount Everest, pounds of meat an American eats per day, year the telephone was invented, and maximum speed of a house cat. Anchoring effect was quantified by the deviation of an estimate from the group mean estimate in the direction of the anchor, normalized to the group estimates’ range. Data from 3 participants whose estimates were clear outliers (i.e., whose distance from others’ estimates was more than ten times the range of others’ estimates) and 1 participant with fewer than two valid (i.e., mostly artifact free) pupil response measurements were excluded from the analysis.

### Persistence of belief experiment

Participants were presented with two urns filled with colored balls (A), and with a sequence of 90 balls, which they were told were sampled with replacement from one of the urns (Peterson & DuCharme, 1967). Every 5 balls, participants indicated using a sliding bar which urn they thought the sequence was sampled from. The precise position of the bar indicated degree of certainty. The first 3 participants performed a preliminary version of the experiment in which they responded after every ball. This was changed to make the experiment faster and more engaging, and thus, 41 participants performed the final version of the experiment. One urn contained 3 red balls, 2 green balls, 2 blue balls, 2 brown balls and 1 purple ball, and the other urn contained 2 red balls, 3 green balls, 1 blue ball, 2 brown balls and 2 purple balls. The sequence of balls was set up so that the first 30 balls favored one of the urns as their source with a probability of 0.95, and the next 60 balls favored the other urn to a similar degree (per 30 balls). Therefore, it was optimal to favor one urn after 30 balls, be indifferent after 60 balls, and favor the other urn after 90 balls (Fig. 10). Accordingly, an optimal observer would be indifferent on average during the last 60 balls. Thus, persistence-of-belief effect was quantified by the degree to which participants’ average response during the last 60 balls favored the initially-favored urn. The initially-favored urn was counterbalanced between participants. Data from 4 participants who did not favor the correct urn during the first 30 balls and 2 participants with fewer than two valid pupil response measurements were excluded from the analysis.

**Fig. 10.**
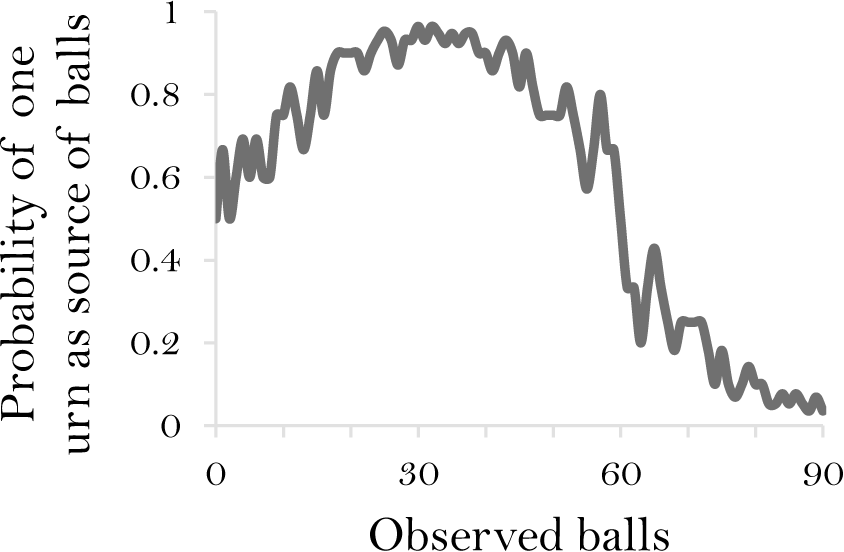
Probability of one urn being the source of the sequence of balls as the sequence progressed, determined by the relative likelihood of each of the balls coming out of the urn, given the contents of both urns.

### Attribute framing experiment

Participants used a sliding bar to rate ground beef products, gambles, and students’ performance, whose attributes were framed either positively or negatively (Levin et al., 1985). In the ground beef task, participants were asked to imagine that they were having a friend over for dinner and they were about to make their favorite lasagna dish with ground beef. They were then asked to rate how satisfied they would be purchasing each of 4 ground beef products, described in terms of price per pound ($2.7 and $3.3), and either percentage lean (80% and 90%, positive frame) or percentage fat (20% and 10%, negative frame). In the gambles task, participants were asked to imagine that they started out with $10 and they can either keep the $10 and not play the gamble or pay the $10 to take the gamble. They were then asked to rate how likely they were to take each of 3 gambles, described in terms of amount to be won ($50, $100 and $200) and either probability of wining (20%, 10% and 5%, positive frame) or probability of losing (80%, 90% and 95%, negative frame). In the student performance task, participants were asked to evaluate each of 2 students on the basis of midterm exam and final exam performance, described in terms of either % correct (50% and 70%, positive frame) or % incorrect (50% and 30%, negative frame). Each item was framed positively in half of the participants, and negatively in the other half. For a given participant, all items of a particular type were similarly framed (i.e., either positively or negatively), so as to minimize awareness of the framing manipulation, but framing was varied within participants across item types. Framing effect was quantified for each item type by the deviation of a participant’s mean rating from the group mean rating in the direction of the frame (i.e., upwards for positive frames, and downwards for negative frames). Data from 1 participant with fewer than two valid pupil response measurements were excluded from the analysis.

### Risky choice framing experiment

Participants faced two different scenarios, a medical scenario and a fire scenario, and indicated using a sliding bar which of two available actions they would choose in each scenario. One action had a certain outcome and the other an uncertain outcome, both of which were framed in terms of either gains or losses. Scenarios were described in full as done previously (Van Schie & Van Der Pligt, 1995). In the medical scenario, which concerned the treatment of a deadly disease at an island inhabited with 600 inhabitants, participants chose between the gain-framed outcomes ‘300 people will be saved’ and’a 50% chance that 600 people will be saved and a 50% chance that none of the people will be saved’, or between the loss-framed outcomes’300 people will die’ and’a 50% chance that 600 people will die and a 50% chance that none of the people will die’. In the fire scenario, which concerned the treatment of fires threatening 9000 acres of forest, participants chose between the gain-framed outcomes’3000 acres of forest will be saved’ and’a 60% chance that 5000 acres will be saved and a 40% chance that no forest under threat will be saved’, or between the loss-framed outcomes’6000 acres of forest will be lost’ and’a 60% chance that 4000 acres will be lost and a 40% chance that 9000 acres will be lost’. Framing effect was quantified as the deviation of a participant’s preferences from the group mean in the direction of the frame (i.e., towards the certain outcome in the gain frame, and towards the uncertain option in loss frame). Data from 2 participants with no valid pupil response measurements were excluded from the analysis.

### Task framing experiment

Participants faced 5 different problems, concerning various subjects such as child custody, vacation choice, ice-cream choice and gambling. Each problem involved one option that had more positive and negative attributes (the ‘enriched’ option) and one option that had fewer positive and negative attributes (the ‘impoverished’ option). In each problem, half of the participants were asked to choose an option, and the other half were asked to reject an option. For example, in one problem participants were asked to imagine that they served on the jury of an only-child sole-custody case following a relatively messy divorce, and they decided to base their decision entirely on the following few observations. Parent A: average income, average health, average working hours, reasonable rapport with the child, relatively stable social life (this parent has no particularly positive or negative attributes). Parent B: above-average income, very close relationship with the child, extremely active social life, lots of work-related travel, minor health problems (this parent has 3 positive and 2 negative attributes). Half of the participants were asked to which parent they would award sole custody of the child, while the other half were asked which parent they would deny sole custody of the child. Framing effect was quantified by the degree to which across tasks (i.e., award and reject) participants preferred the enriched option (i.e., Parent A) more frequently than the impoverished option (i.e., Parent B). Full description of the other problems can be found elsewhere (Shafir, 1993; problems 1, 2, 4, 5 and 6). Data from 2 participants with fewer than two valid pupil response measurements were excluded from the analysis.

### Sample-size neglect experiment

Participants were told to imagine that they were spinning a biased coin and recording how often the coin landed heads and how often the coin landed tails. They knew that the coin tended to land on one side 3 out of 5 times, but they did not know if this bias is in favor of heads or in favor of tails. Participants were then presented with 10 different sets of results (number of heads and number of tails), in which the heads always outnumbered the tails, and they indicated using a sliding bar how certain they were given each set that the coin was biased in favor of heads. Sets of results were similar to those used previously (Griffin & Tversky, 1992).

As shown by Griffin & Tversky (1992), the probability that the coin was biased in favor of heads according to Bayes’ rule is:

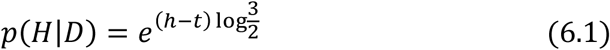

where *h* is the number of heads and *t* is the number of tails. This expression is equivalent to

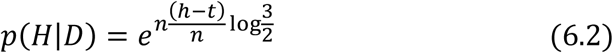

which depends on the sample size (i.e., the number of outcomes, *n*) and on the ratios of heads
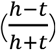
,It was previously shown that people tend to overweigh the ratio component at the expense of the sample size component (sample-size neglect; Griffin & Tversky, 1992). To measure this bias, we regressed participants’ estimates against the true probabilities (Eq. 6.1), and then regressed the residuals against the ratio component alone 
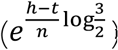
, to measure the degree of overweighting of the ratio component. In addition, to test whether the difference between heads and tails explained participants’ estimates better than the ratio between them, we reversed the steps. That is, we first regressed participants’ estimates against the ratio component, and then the residuals against the true probabilities, which reflect the difference between heads and tails (Eq. 6.1). We then compared the resulting regression coefficients to the coefficients produced when the regressions were performed in the reverse order. All inputs to regression analyses were *z* scored so as to produce normalized coefficients. 7 participants who were more certain that the coin was biased in favor of heads given 3 heads and 2 tails, than given 7 heads and 2 tails, were excluded from the analysis, as we suspected that they mistakenly looked for a ratio that best matched 3 to 2. Including these participants in the analysis did not affect the observed significant relationship between pupil dilation and sample-size neglect in the task.

### Eye tracking

A desk-mounted SMI RED 120Hz eye-tracker (SensoMotoric Instruments Inc., MA) was used to measure participants’ left and right pupil diameters at a rate of 60 samples per second while they were performing the behavioral tasks with their head fixed on a chinrest. At the beginning of the experiment, a baseline measurement of pupil diameter at rest was taken for a period of 45 s. Pupil-diameter data were processed in MATLAB to detect and remove blinks and other artifacts. For each trial, baseline pupil diameter was computed as the average diameter over a period of 1 s prior to the beginning of the trial (at the end of the inter-trial interval, at which point pupil activity from the trial itself should have subsided). Pupil-dilation response was computed as the difference between the peak diameter recorded during the 4 s that followed the beginning of the trial and the preceding baseline diameter. All pupil dilation responses were normalized by the pre-experiment baseline pupil diameter. Pupil dilation responses in which more than half of the measurements were affected by artifacts were considered invalid and excluded from the analysis.

### Statistical analysis

Analyses were carried out using MATLAB. Permutation tests were performed by sampling 100,000 random permutations of the coupling between pupillary and behavioral individual data sets. Results based on the permuted data served as null distributions to which actual results were compared. Since effect sizes approached zero in multiple participants and thus could not be expected to follow a linear relationship, correlations involving effect sizes were computed using Spearman’s rank correlation. Averaging of correlation coefficients was preceded by Fisher’s transformation so as to mitigate the problem of the non-additivity of correlation coefficients (Fisher, 1921). All statistical tests were two tailed.

### Computational model

We modeled decision between two items, each with two attributes, using a leaky competing accumulator model (Usher & McClelland, 2001). The model consisted of two competing accumulators (A), one for each item. Every timestep, activity ai of accumulator i was updated to reflect evidence in favor of the respective item by:

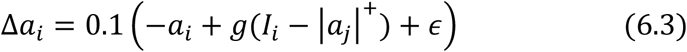

where *g* reflects the level of gain, *I*_i_ is the evidence-based excitatory input to accumulator *i*, *j* is the competing accumulator whose positive component (||+) provided inhibitory input, and *є* is zero-mean normally-distributed noise with a standard deviation of 0.01. As in previous models of multi-attribute multi-item decisions (Usher & McClelland, 2004), on each timestep, one attribute was selected at random, and excitatory input was determined accordingly. One of the attributes favored one item, and thus generated input of 1.25 to one accumulator and 0.75 to the other accumulator (plus normally-distributed random noise with standard deviation of 0.01). The other attribute favored the other item, and thus generated the same input but reversed. A decision was reached once one of the accumulators reached a value of 1. A bias was implemented by either setting the likelihood of selecting one of the attributes to 0.55 (instead of 0.5), or by increasing the input to one of the accumulators by 0.05. The two implementations gave similar results, and thus only the results of the first are shown. We conducted 1,000,000 simulations with each level of gain between 1 and 10.

## Acknowledgements

This project was made possible through a grant from the Howard Hughes Medical Institute to EE.

## References

Aston-Jones G, Cohen, JD (2005) An integrative theory of locus coeruleus-norepinephrine function: adaptive gain and optimal performance. Annu Rev Neurosci 28:403–450 (2005).

Busemeyer JR, Jessup RK, Johnson JG, Townsend JT (2006) Building bridges between neural models and complex decision making behaviour. Neural Networks 19(8):1047–1058.

Busemeyer JR, Townsend JT (1993) Decision field theory: a dynamic-cognitive approach to decision making in an uncertain environment. Psychol Rev 100(3):432.

Daw ND, Niv Y, Dayan P (2005) Uncertainty-based competition between prefrontal and dorsolateral striatal systems for behavioral control. Nat Neurosci 8(12):1704–1711.

Diederich A (1997) Dynamic stochastic models for decision making under time constraints. J Math Psychol 41(3):260–274.

Eldar E, Cohen JD, Niv Y (2013) The effects of neural gain on attention and learning. Nat Neurosci 16(8):1146–1153.

Eldar E, Niv Y, Cohen JD (submitted) Neural gain and integration in perceptual processing.

Glöckner A, Betsch T (2008) Modeling option and strategy choices with connectionist networks: Towards an integrative model of automatic and deliberate decision making. Judgm Decis Mak 3:215–228.

Griffin D, Tversky A (1992) The weighing of evidence and the determinants of confidence. Cogn Psychol 24(3):411–435.

Jacowitz KE, Kahneman D (1995) Measures of anchoring in estimation tasks. Pers Soc Psychol Bull 21:1161–1166.

Johnson JG, Busemeyer JR (2005) A dynamic, stochastic, computational model of preference reversal phenomena. Psychol Rev 112(4):841.

Joshi S, Li Y,Kalwani RM, Gold JI (2016) Relationships between pupil diameter and neuronal activity in the locus coeruleus, colliculi, and cingulate cortex. Neuron 89(1), 221–234.

Kahneman D (1973)Attention and effort (Prentice-Hall, Englewood Cliffs, NJ).

Kahneman D, Tversky A (1979) Prospect theory: An analysis of decision under risk. Econometrica 47(2):363–391.

Krajbich I, Rangel AA (2011) multi-alternative drift diffusion model predicts the relationship between visual fixations and choice in value-based decisions. Proc Natl Acad Sci USA 108(33):13852–13857.

Kühberger A (1998) The influence of framing on risky decisions: A meta-analysis. Organ Behav Hum Decis Process 75(1):23–55.

Levin IP, Johnson RD, Russo CP, Deldin PJ (1985) Framing effects in judgment tasks with varying amounts of information. Organ Behav Hum Decis Process 36(3):362–377.

Levin IP, Schneider SL, Gaeth GJ (1998) All frames are not created equal: A typology and critical analysis of framing effects. Organ Behav Hum Decis Process 76(2):149–188.

Lord CG,Ross L, Lepper MR (1979) Biased assimilation and attitude polarization: The effects of prior theories on subsequently considered evidence. J Pers Soc Psychol 37(11):2098.

Peterson CR, DuCharme WM (1967) A primacy effect in subjective probability revision. J Exp Psychol 73(1):61.

Roe RM, Busemeyer JR, Townsend JT (2001) Multialternative decision field theory: A dynamic connectionst model of decision making. Psychol Rev 108(2):370.

Rusou Z, Zakay D, Usher M (2013) Pitting intuitive and analytical thinking against each other: The case of transitivity. Psychon Bull Rev 20(3):608–14.

Servan-Schreiber D, Printz H, Cohen JD (1990) A network model of catecholamine effects: gain, signal-to-noise ratio, and behavior. Science 249(4971):892–895.

Shafir E (1993) Choosing versus rejecting: Why some options are both better and worse than others. Mem Cognit 21(4):546–556.

Tversky A, Kahneman D (1974) Judgment under uncertainty: Heuristics and biases. Science 185(4157):1124–1131.

Tversky A, Kahneman D (1981) The framing of decisions and the psychology of choice. Science 211(4481):453–458.

Usher M, McClelland JL(2004) Loss aversion and inhibition in dynamical models of multialternative choice. Psychol Rev 111(3):757.

Usher M, Russo Z, Weyers M, Brauner R, Zakay D (2011) The impact of the mode of thought in complex decisions: Intuitive decisions are better. Front Psychol 2:37.

Usher M, Tsetsos K, Erica CY, Lagnado DA (2013) Dynamics of decision-making: from evidence accumulation to preference and belief. Front Psychol 4:758.

Van Schie EC, Van Der Pligt J (1995) Influencing risk preference in decision making: The effects of framing and salience. Organ Behav Hum Decis Process 63(3):264–275.

